# An interventional Soylent diet increases the *Bacteroidetes* to *Firmicutes* ratio in human gut microbiome communities: a randomized controlled trial

**DOI:** 10.1101/200881

**Authors:** Ryan H. Hsu, Dylan M. McCormick, Mitchell J. Seitz, Lauren M. Lui, Harneet S. Rishi, Adam P. Arkin

**Author notes:** Corresponding author: (RH).

## Abstract

Our knowledge of the relationship between the gut microbiome and health has rapidly expanded in recent years. Diet has been shown to have causative effects on microbiome composition, which can have subsequent implications on health. Soylent 2.0 is a liquid meal replacement drink that satisfies nearly 20% of all recommended daily intakes per serving. This study aims to characterize the changes in gut microbiota composition resulting from a short-term Soylent diet. Fourteen participants were separated into two groups: 5 in the regular diet group and 9 in the Soylent diet group. The regular diet group maintained a diet closely resembling self-reported regular diets. The Soylent diet group underwent three dietary phases: A) a regular diet for 2 days, B) a Soylent-only diet (five servings of Soylent daily and water as needed) for 4 days, and C) a regular diet for 4 days. Daily logs self-reporting diet, Bristol stool ratings, and any abdominal discomfort were electronically submitted. Eight fecal samples per participant were collected using fecal sampling kits, which were subsequently sent to uBiome, Inc. for sample processing and V4 16S rDNA sequencing. Reads were clustered into operational taxonomic units (OTUs) and taxonomically identified against the GreenGenes 16S database. We find that an individual’s alpha-diversity is not significantly altered during a Soylent-only diet. In addition, principal coordinate analysis using the unweighted UniFrac distance metric shows samples cluster strongly by individual and not by dietary phase. Among Soylent dieters, we find a significant increase in the ratio of *Bacteroidetes* to *Firmicutes* abundance, which is associated with several positive health outcomes, including reduced risks of obesity and intestinal inflammation.

## Introduction

The gut microbiota plays a key role in mediating human health, including aspects of infection, inflammation, and obesity [1–3]. While many factors that influence the gut microbiome have been identified, diet has been consistently shown to be a major contributor [4,5]. This has led to a growing consumer awareness of dietary choices that function as prebiotics and probiotics [6].

Meal replacement drinks are an emerging alternative to traditionally prepared meals that are designed to conveniently supply full servings of dietary nutrients. Among these products, Soylent 2.0 in particular has gained recent popularity. Soylent 2.0’s formulation is engineered to fulfill nearly all recommended daily intakes of total fat, sodium, potassium, carbohydrates, proteins, vitamins, and minerals, without excess sugars or cholesterol [7]. Given the critical role of intestinal flora in various human health outcomes, we are interested in studying how meal replacements like Soylent affect gut microbiome composition.

To assess the compositional changes to the gut microbiome resulting from a Soylent-only diet, we performed a parallel randomized controlled trial consisting of a control group that adhered to self-reported regular diets and a treatment group that received a Soylent 2.0 dietary intervention. Fecal samples were regularly collected and sequenced to quantify microbial populations at defined timepoints throughout the study.

This study was conceived, designed, and coordinated by a team of undergraduates at UC Berkeley and funded via Experiment, a website that hosts online crowdfunding campaigns for scientific research. The funders did not participate in study conception, experimental design, or data analysis.

## Material and Methods

### Institutional clinical trial registration

The study was granted institutional review board approval from the Committee for Protection of Human Subjects at the University of California, Berkeley (CPHS #2016-04-8727, October 14, 2016). The trial was publicly submitted to the clinicaltrials.gov registry (ID #NCT03203044, June 27, 2017) following completion. The trial was not publicly registered prior to subject enrollment due to a miscategorization of the study as a non-clinical trial. All related and future trials will be prospectively submitted to a public registry.

### Subject enrollment and selection

Written informed consent to participate in the study was solicited through Mycrobes, an undergraduate student organization at the University of California, Berkeley. Consenting individuals were administered a questionnaire surveying for age, biological sex, ethnicity, student status, pregnancy status, food allergies or sensitivities, health complications, dietary supplements or medications, prior Soylent consumption, and dietary and lifestyle descriptions. Participants were selected at random from eligible individuals who did not report non-student status, pregnancy, serious food allergies or sensitivities, chronic disease, current use of medications, or prior Soylent consumption (Fig 1). Participation in the study was entirely voluntary, and subjects did not receive any compensation.

**Fig 1.**
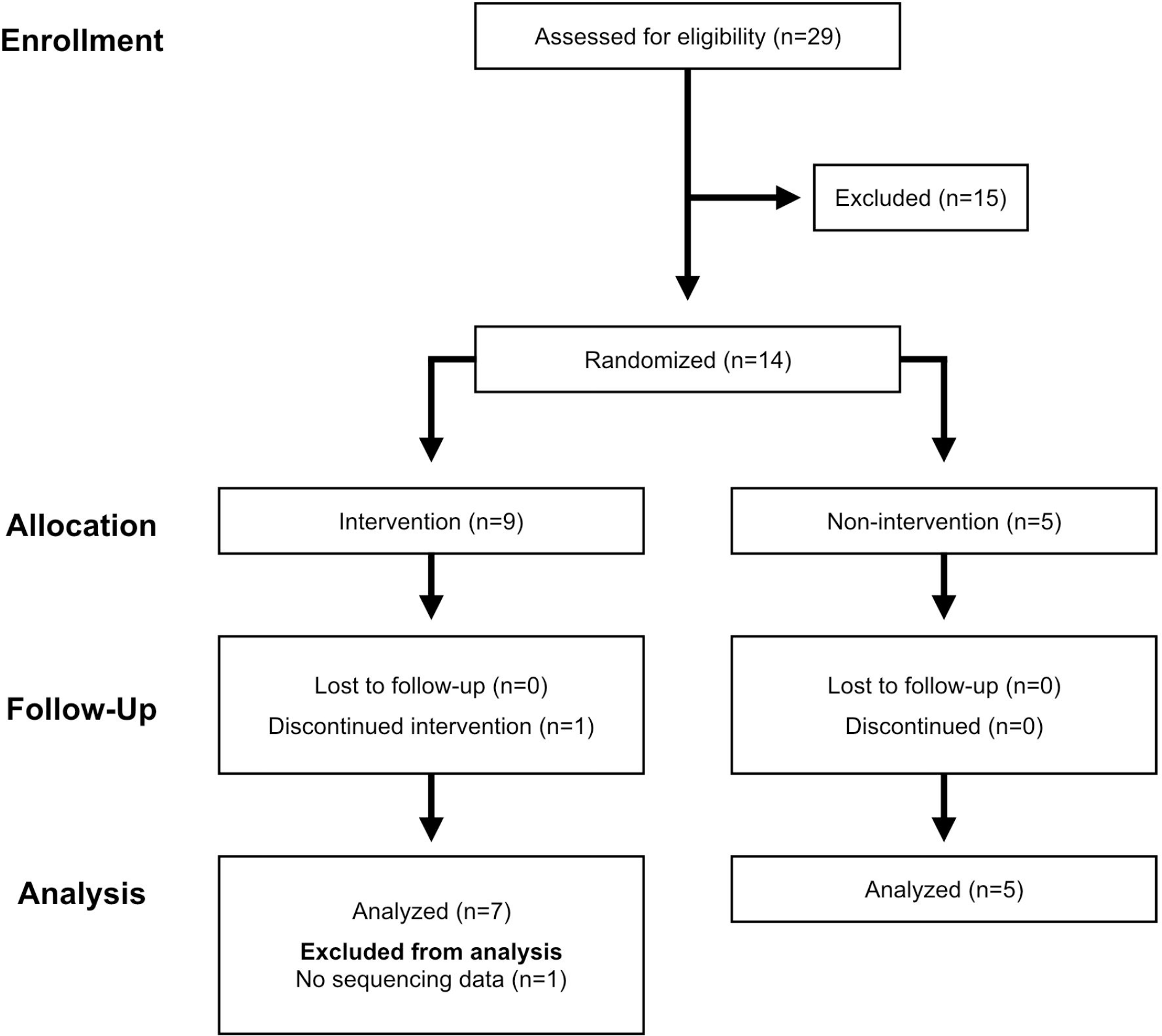
CONSORT flow diagram of the study.

### Study procedure

The study design is a parallel randomized controlled trial. Participants were randomly assigned to the control or treatment group, also referred to as the regular diet and Soylent diet group. Individuals in the control group maintained their regular diets throughout the study, allowing characterization of daily fluctuations in microbiome composition, while participants in the treatment group received 20 bottles of Soylent 2.0 to consume during phase B of the study. Soylent 2.0 is a liquid formulation consisting of primarily soy protein, algal oil, and isomaltulose, as well as smaller amounts of other ingredients such as essential vitamins and minerals (S1 Fig).

The study was organized into three phases spanning a period of ten days (Fig 2). During phase A, all participants maintained their regular diets for two days. In phase B, the Soylent diet group switched to a Soylent-only diet consisting of a recommended 5 servings of Soylent daily *ad libitum* (and water as needed) for four days, while the regular diet group retained a normal diet. During Phase C, all participants returned to their regular diets for four days. Prior to the initiation of the study, eight uBiome stool sampling kits were distributed to each participant for use on eight specific days. Participants who missed a sampling day were instructed to sample on the following day if possible. Additionally, participants submitted daily electronic forms reporting their diet, time of bowel movements, Bristol stool ratings, and any symptoms or discomfort (S1 File). The primary outcome measure is microbiome composition.

**Fig 2.**
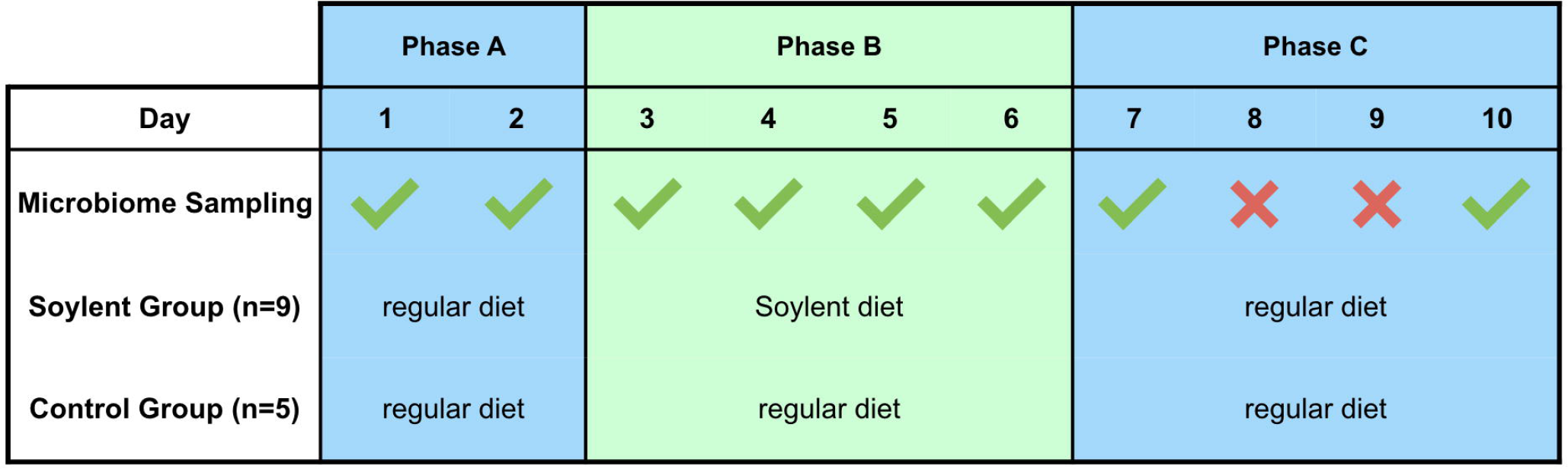
Study design. The study design consists of a time-course over 10 days. 9 and 5 participants were randomized to the Soylent and regular diet groups, respectively. The Soylent diet group maintained a regular diet during phase A to quantify baseline microbiome composition, switched to a Soylent-only diet during phase B, and returned to a regular diet during phase C. The regular diet group, which serves a control, maintained a regular diet throughout the study period to quantify day-to-day variations in microbiome composition. Fecal samples were collected on the 8 days specified using the uBiome Gut Kit and submitted for 16S sequencing.

### Sample collection and 16S rDNA sequencing

Commercially available uBiome Gut Kits were used to sample, store, and transport fecal samples to uBiome, Inc [8]. Fecal samples were swabbed from used toilet paper with the included sterile swabs and immediately resuspended into tubes containing lysis and stabilization buffer. The tubes were sealed and stored at ambient temperature. Once all sampling was completed, samples were delivered to uBiome Inc. and run through their standard stool 16S sample processing and sequencing pipeline as follows: Genomic DNA was extracted in a class 1000 clean room using a column-based approach by a liquid-handling robot. The V4 region of the 16S rRNA gene was amplified using universal primers (515F:GTGCCAGCMGCCGCGGTAA and 806R:105 GGACTACHVGGGTWTCTAAT) and barcoded to allow for multiplexed sequencing. Column-based cleanup, size selection, and qPCR quantification were utilized to prepare libraries. 2 × 150 bp paired-end sequencing was performed on an Illumina NextSeq 500. Sequencing reads were demultiplexed using the Illumina BCL2FASTQ algorithm and electronically sent to the researchers.

### Read processing and OTU analysis

Custom scripts were used to merge sequencing data across sequencing lanes and relabel samples [9]. USEARCH 9.2 was utilized to stitch and quality filter the reads. Chimeras were identified de-novo and removed with the UCHIME algorithm [10,11]. Reads were then clustered and assigned to OTUs at a 0.97 identity threshold using VSEARCH 2.3.4 [12]. Mothur 1.39.5 was run to assign taxonomy according to the GreenGenes_13_8 16S database, allowing the determination of abundance at near-species resolution in each sample [13,14]. PyNAST 1.2.2 and FastTree were employed to align OTUs and build a tree used to infer phylogenetic distance [15,16]. Shannon-Wiener and Gini-Simpson ecological diversity indices and UniFrac distance metrics were calculated using QIIME 1.9.1 [17]. To reduce biases arising from varying sequencing depth, samples with less than 2,000 reads were discarded (n=6), and the remaining samples (n=62) were normalized by rarefying to 2,000 sequences [18]. Rarefaction curves and jackknife estimates were generated to validate the 2,000 read threshold (S2 File) [17]. Samples were binned into one of six groups representing the diet arm (Regular or Soylent) and study phase (A, B, or C). Since it has been shown that stomach contents take about a day to fully reach and pass the intestinal tract, samples are categorized by the dietary phase 24 hours prior to collection [6].

### Taxonomic analysis

Intra-sample diversity (α-diversity) was measured using the Shannon-Wiener (H) and Gini-Simpson (D) diversity indices [17]. Calculated α-diversity metrics were averaged within one of six bins representing diet-group and phase of study (e.g. Soylent A). For each individual, the relative abundances of four dominant phyla (*Actinobacteria*, *Bacteroidetes*, *Firmicutes*, and *Proteobacteria)* were normalized by subtracting the baseline community as established in phase A. These values were then binned and averaged in the same manner as the aforementioned α-diversity metrics.

Distributions within bins containing at least eight samples (Soylent A, Soylent B, Regular A, and Regular B) were tested for normality using the D’Agostino-Pearson omnibus test. For α-diversity metrics, the Regular A, Soylent A, and Soylent B bins were accepted to follow normal distributions (*p* > 0.05), while the Regular B bin was not (*p* < 0. 05). For changes in the relative abundance of *Bacteroidetes*, *Firmicutes*, and *Proteobacteria*, the Regular A, Regular B, Soylent A, and Soylent B bins were accepted to follow normal distributions (*p* > 0.05). For changes in the relative abundance of *Actinobacteria*, the Soylent A and Soylent B bins were accepted to follow normal distributions (*p* > 0.05), while the Regular A and Regular B bins were not (*p* < 0.05).

Differences between these bins were statistically analyzed using two-tailed t-tests, and *p*-values less than 0.05 were considered to be statistically significant.

## Results

### Selected cohort

Study recruitment and follow-ups began on October 20, 2016 and ended on December 2, 2016. Informed Consent was received from 29 individuals, and screening forms were subsequently collected to determine which individuals were eligible for participation. Fourteen participants were randomized into either the regular diet or Soylent diet group with a 5:9 allocation ratio using the ‘random.permutation’ function of the NumPy package for Python 2.7 (Fig 1). The sample size was determined to maximize use of study funding.

The 14 participants were all active University of California, Berkeley undergraduates aged 18-21. The regular diet group consisted of 2 females and 3 males, while the Soylent diet group consisted of 3 females and 6 males. Participants were of Asian, European, Hispanic, and Native American descent. Three participants identified as vegetarians, while the rest consumed vegetables and meat on a regular basis (Table 1).

**Table 1. Baseline sociodemographic and dietary profiles of the participants.**

### Quality of collected samples

One individual in the Soylent diet group consumed less Soylent than instructed, and therefore their data was not included in the analysis. Eight of the remaining 104 fecal samples were not collected due to a lack of bowel movements. Of the submitted samples, 71.9% returned sequencing data from uBiome Inc., with the 1st, 2nd, and 3rd quartiles of sequencing depths at 8335, 83777, and 208145 reads, respectively (S1 Table, S2 Fig). Although we also collected Bristol stool ratings from each participant, the data were inconsistent and did not warrant formal analysis.

### α- and β-diversity metrics remain stable during a Soylent diet

Shannon-Wiener and Gini-Simpson α-diversity indices did not significantly change from a Soylent dietary intervention (phase A to B) (S3 Fig).

Inter-sample diversity (β-diversity) was quantified using unweighted and weighted UniFrac, which scores distance between samples according to phylogenetic similarity, in this case using the 16S V4 sequence. β-diversity was visualized using principal-coordinate analysis (PCoA). Unweighted Unifrac, which only considers the presence or absence of each OTU, shows that samples strongly cluster by individual. No clustering of Soylent-treated samples (those taken from participants in the Soylent diet group during phase B) is observed (Fig 3). Similar clustering patterns are visible using weighted UniFrac, which scores similarity of samples based on OTU abundance as well (S4 Fig).

**Fig 3.**
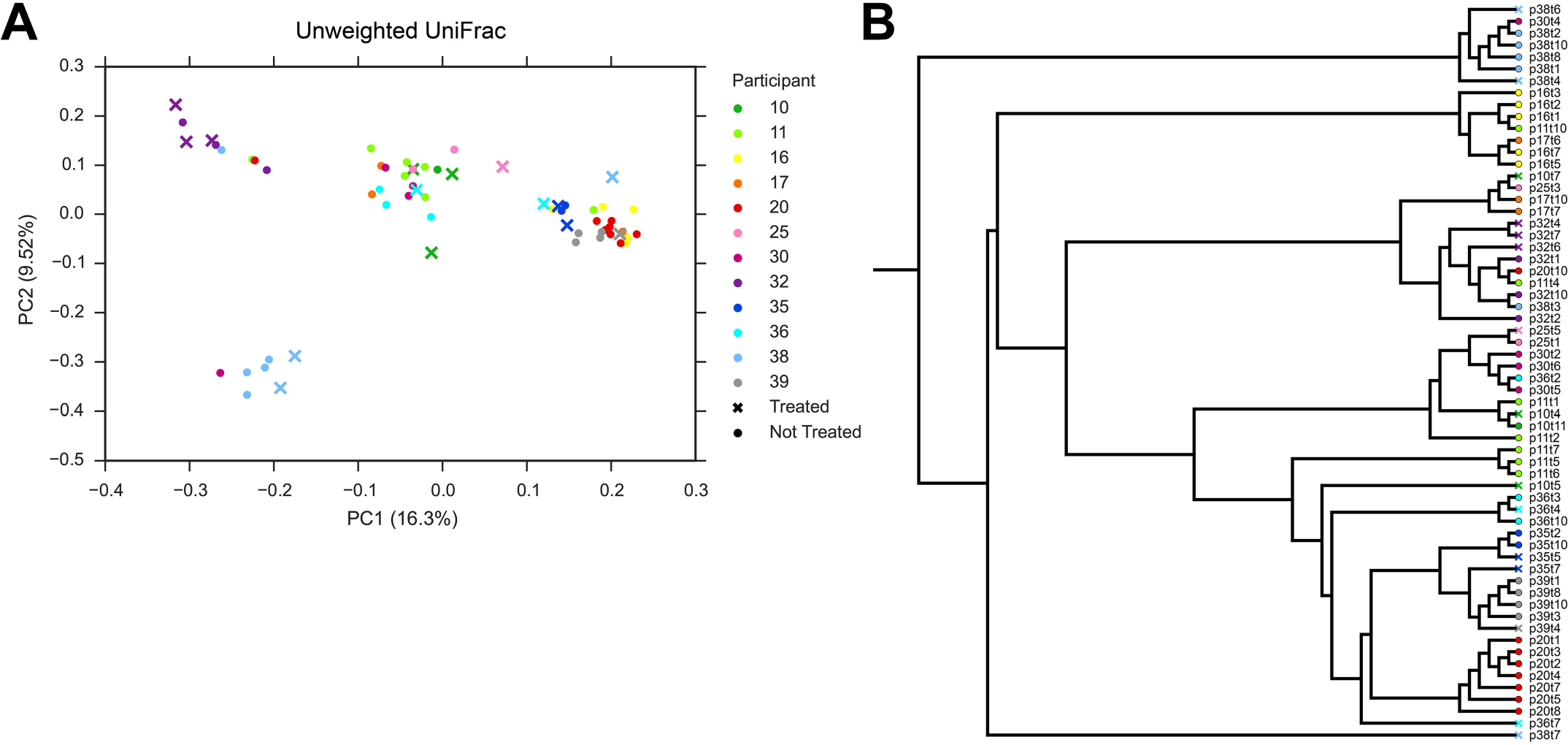
Principle coordinate analysis and visualization of the unweighted UniFrac metric. A) Unweighted UniFrac, which considers the binary presence or absence of each OTU, reveals that samples cluster strongly by participant (indicated by color) and not by diet (denoted by •’s or ×’s). B) Agglomerative clustering performed on the first ten principal coordinates validates these clustering patterns.

### Soylent consumption alters relative abundance of dominant phyla

During the Soylent dietary intervention (phase B), the Soylent diet group exhibited a significant increase in the abundance of *Bacteroidetes* (*p*=0.011), as well as a statistically insignificant decrease in the abundance of *Firmicutes* (*p*=0.078), compared to the regular diet group (Fig 4A). Accordingly, the *Bacteroidetes* to *Firmicutes* ratio increased significantly during the same time period (*p*=0.028) (Fig 4B). No significant change in *Proteobacteria* abundance was observed (Fig 4A). Since the *Actinobacteria* bins did not follow normal distributions, no comparison was made for the phylum.

**Fig 4.**
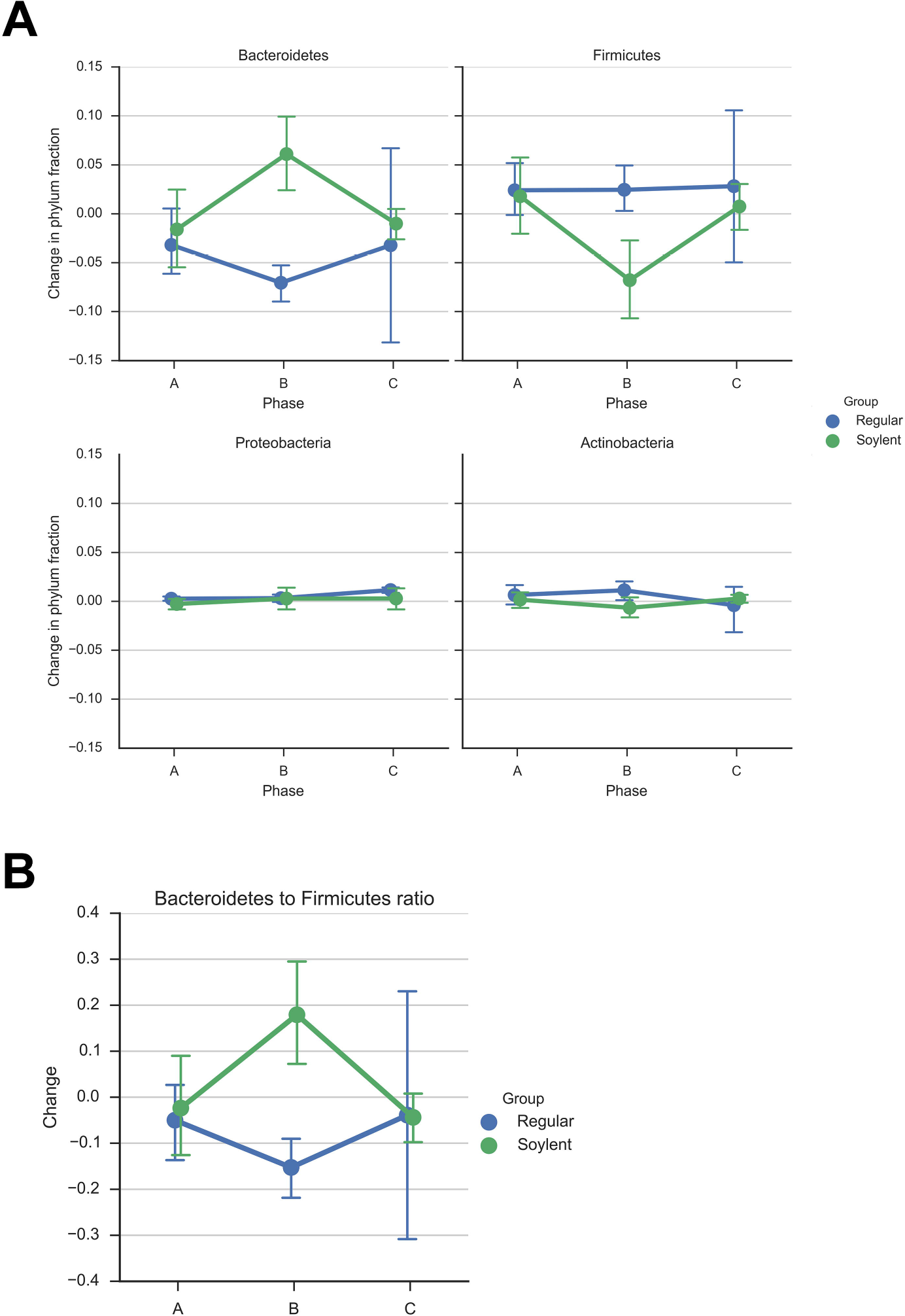
Abundance changes of dominant gut microbiota phyla. A) Changes in the normalized relative abundance of four dominant gut microbiota phyla averaged within each diet-phase group. In comparison with the regular diet group, the Soylent diet group shows a significant increase in *Bacteriodetes* (*p*=0.011) and an insignificant decrease in *Firmicutes* (*p*=0.078) abundances during the Soylent dietary intervention (phase B). There is no significant change in *Proteobacteria* abundance during this phase (*p*=0.937). Statistical comparison of *Actinobacteria* abundance was not performed because the data was not normally distributed. B) Changes in the *Bacteriodetes* to *Firmicutes* ratio. During the Soylent diet intervention (phase B), there is a significant increase in the *Bacteroidetes* to *Firmicutes* ratio (p=0.028). Standard error is shown.

### Discussion

Based on calculated Shannon-Wiener and Gini-Simpson diversity scores, we find no significant change in α-diversity across either diet arm, which suggests that overall microbiome diversity is resilient to dietary changes in the short term. Since Soylent is very nutrient-rich, it likely does not starve a large enough fraction of gut microbiota to significantly reduce α-diversity. This finding could also be by the ability of gut microbes to remain dormant during the Soylent dietary intervention. Microbiome diversity is of clinical significance and is negatively associated with diseases such as recurrent *Clostridium difficile* infection [19]. We find that the interventional Soylent diet does not negatively impact gut microbiota diversity.

Unweighted UniFrac reveals that samples cluster strongly within an individual. Furthermore, samples from Soylent diet days among different participants do not cluster together, but rather cluster strongly with each individual’s other samples. Visualization of weighted UniFrac, which scores sample similarity based on OTU abundance as well, exhibits similar clustering patterns. The clustering patterns described are less apparent, possibly due to the steep rarefaction that was performed to normalize sample read depth. This suggests that Soylent does not significantly change which organisms are present in the microbiome, which can be partially explained by the fact that Soylent is pasteurized and therefore cannot act as a probiotic.

We find that Soylent consumption significantly increases the *Bacteroidetes* to *Firmicutes* ratio in the gut microbiota. Since the *Bacteroidetes* and *Firmicutes* together compose over 90% of the sampled microbiomes, their abundances are expected to be inversely correlated. An increase in this ratio is observed in phase B of the Soylent diet arm compared to the regular diet arm, followed by a rapid return to baseline levels in phase C. Previous studies have demonstrated associations between the relative abundance of the phyla *Bacteroidetes* and *Firmicutes* and specific health outcomes. In particular, it has been shown that a low *Bacteroidetes* to *Firmicutes* ratio is associated with obesity in mice and humans [20,21]. Additionally, a low ratio is associated with ulcerative colitis and Crohn’s disease, which are types of inflammatory bowel disease [22]. Although these associations have been strongly demonstrated, causality has not yet been established in either case.

Although the participant pool is relatively small, similar sample sizes have articulated clear results in other diet-related gut microbiome studies [6]. Nevertheless, missing data for some samples presents a clear limitation to the study. We were unable to troubleshoot these samples because sequencing was outsourced to uBiome Inc. Additionally, the number of reads per sample was highly variable. Therefore, we had to rarefy our samples to perform meaningful statistical analyses. Although we discarded many reads from high-depth samples, rarefying has been shown to effectively mitigate biases associated with sequencing depth variation [18].

Although the limitations of 16S surveys are well understood (particularly the lack of access to functional information), the accuracy of metagenome reconstruction is still under debate and conclusions drawn from this method should be met with skepticism. To confidently address these limitations, further studies involving transcriptomics and metagenomics must be conducted to more accurately identify changes in gene expression, gene function, and abundance resulting from a Soylent 2.0 dietary intervention.

We conclude that a short-term interventional Soylent diet does not negatively impact the composition of gut flora communities. As additional studies demonstrate the effect that specific microbial consortia have on health, both food product manufacturers and consumers should consider the gut microbiome as an essential component of human health.

## Data Availability

Raw sequencing reads in FASTQ format from this study were uploaded to EBI’s European Nucleotide Archive under the accession code PRJEB21752 (http://www.ebi.ac.uk/ena/data/view/PRJEB21752).

## Acknowledgements

We greatly appreciate the 84 backers of our Experiment crowdfunding campaign for making this study possible. We thank Gwyneth Terry for assisting with the administrative coordination of the project, as well as Shyam Bhakta and Michael Shapira for helpful discussions. We also thank uBiome Inc. for their partnership in performing sample processing and DNA sequencing.

## Supporting Information

**S1 Fig. Nutrition Facts for the Soylent 2.0 meal replacement drink.** Participants in the Soylent diet group consumed 5 servings of Soylent per day during phase B of the study.

**S2 Fig. Sequencing depth was variable among samples.** The 1st, 2nd, and 3rd quartiles of sequencing depths were 8335, 83777, and 208145 reads, respectively.

**S3 Fig. α-diversity remains consistent throughout a Soylent dietary intervention.** The Soylent dietary intervention did not result in a significant change in α-diversity (Soylent A to Soylent B), as shown by the Shannon-Wiener (*p*=0.279) and Gini-Simpson (*p*=0.999) metrics. Standard error is shown.

**S4 Fig. Principle Coordinate Analysis visualization of two Unifrac variants.** Unweighted UniFrac, which considers the binary presence or absence of each OTU, reveals that samples cluster strongly by participant (indicated by color) and not by diet (denoted by •’s or ×’s). Similar but less apparent trends are seen with the Weighted UniFrac metric, which weights OTUs based on fractional abundance as well.

**S1 Table.** The table links participant IDs to diet groups and collected stool samples. Samples that returned no reads or were discarded due to low sequencing depth are denoted.

**S1 File. Daily Electronic Log Form**

**S2 File. Validation and discussion of rarifying**

## References

1. Turnbaugh PJ, Ley RE, Mahowald MA, Magrini V, Mardis ER, Gordon JI. An obesity-associated gut microbiome with increased capacity for energy harvest. Nature. 2006;444: 1027–1031.

2. Halfvarson J, Brislawn CJ, Lamendella R, Vázquez-Baeza Y, Walters WA, Bramer LM, et al. Dynamics of the human gut microbiome in inflammatory bowel disease. Nat Microbiol. 2017;2: 17004.

3. Brandt LJ, Aroniadis OC, Mellow M, Kanatzar A, Kelly C, Park T, et al. Long-term follow-up of colonoscopic fecal microbiota transplant for recurrent Clostridium difficile infection. Am J Gastroenterol. 2012;107: 1079–1087.

4. Zhernakova A, Kurilshikov A, Bonder MJ, Tigchelaar EF, Schirmer M, Vatanen T, et al. Population-based metagenomics analysis reveals markers for gut microbiome composition and diversity. Science. 2016;352: 565–569.

5. Smits SA, Marcobal A, Higginbottom S, Sonnenburg JL, Kashyap PC. Individualized Responses of Gut Microbiota to Dietary Intervention Modeled in Humanized Mice. mSystems. 2016;1. doi:10.1128/mSystems.00098-16

6. David LA, Maurice CF, Carmody RN, Gootenberg DB, Button JE, Wolfe BE, et al. Diet rapidly and reproducibly alters the human gut microbiome. Nature. 2014;505: 559–563.

7. Rosa Foods I. Soylent — Food, intelligently designed [Internet]. [cited 4 Jun 2017]. Available: http://soylent.com

8. Almonacid DE, Kraal L, Ossandon FJ, Budovskaya YV, Cardenas JP, Bik EM, et al. 16S rRNA gene sequencing and healthy reference ranges for 28 clinically relevant microbial taxa from the human gut microbiome. PLoS One. 2017;12: e0176555.

9. ryanusahk/mycrobes_soylent_microbiome: Mycrobes Soylent Microbiome DNA Analysis Pipeline. 2017; doi:10.5281/zenodo.822861

10. Edgar RC. Search and clustering orders of magnitude faster than BLAST. Bioinformatics. 2010;26: 2460–2461.

11. Edgar RC, Haas BJ, Clemente JC, Quince C, Knight R. UCHIME improves sensitivity and speed of chimera detection. Bioinformatics. 2011;27: 2194–2200.

12. Rognes T, Flouri T, Nichols B, Quince C, Mahé F. VSEARCH: a versatile open source tool for metagenomics. PeerJ. 2016;4: e2584.

13. Schloss PD, Westcott SL, Ryabin T, Hall JR, Hartmann M, Hollister EB, et al. Introducing mothur: open-source, platform-independent, community-supported software for describing and comparing microbial communities. Appl Environ Microbiol. 2009;75: 7537–7541.

14. McDonald D, Price MN, Goodrich J, Nawrocki EP, DeSantis TZ, Probst A, et al. An improved Greengenes taxonomy with explicit ranks for ecological and evolutionary analyses of bacteria and archaea. ISME J. 2012;6: 610–618.

15. Caporaso JG, Bittinger K, Bushman FD, DeSantis TZ, Andersen GL, Knight R. PyNAST: a flexible tool for aligning sequences to a template alignment. Bioinformatics. 2010;26: 266–267.

16. Price MN, Dehal PS, Arkin AP. FastTree 2--approximately maximum-likelihood trees for large alignments. PLoS One. 2010;5: e9490.

17. Caporaso JG, Kuczynski J, Stombaugh J, Bittinger K, Bushman FD, Costello EK, et al. QIIME allows analysis of high-throughput community sequencing data. Nat Methods. 2010;7: 335–336.

18. Weiss S, Xu ZZ, Peddada S, Amir A, Bittinger K, Gonzalez A, et al. Normalization and microbial differential abundance strategies depend upon data characteristics. Microbiome. BioMed Central; 2017;5: 27.

19. Antharam VC, Li EC, Ishmael A, Sharma A, Mai V, Rand KH, et al. Intestinal Dysbiosis and Depletion of Butyrogenic Bacteria in Clostridium difficile Infection and Nosocomial Diarrhea. J Clin Microbiol. 2013;51: 2884–2892.

20. Ley RE, Bäckhed F, Turnbaugh P, Lozupone CA, Knight RD, Gordon JI. Obesity alters gut microbial ecology. Proc Natl Acad Sci U S A. 2005;102: 11070–11075.

21. Ley RE, Turnbaugh PJ, Klein S, Gordon JI. Microbial ecology: human gut microbes associated with obesity. Nature. 2006;444: 1022–1023.

22. Zhou Y, Zhi F. Lower Level of Bacteroides in the Gut Microbiota Is Associated with Inflammatory Bowel Disease: A Meta-Analysis. Biomed Res Int. 2016;2016: 5828959.

